# Morphological integration during postnatal ontogeny: implications for evolutionary biology

**DOI:** 10.1101/2021.07.13.452198

**Authors:** Alex Hubbe, Fabio A. Machado, Diogo Melo, Guilherme Garcia, Harley Sebastião, Arthur Porto, James Cheverud, Gabriel Marroig

## Abstract

How covariance patterns of phenotypes change during development is fundamental for a broader understanding of evolution. There is compelling evidence that mammalian skull covariance patterns change during ontogeny. However, it is unclear to what extent variation in covariance patterns during ontogeny can impact the response to selection. To tackle this question we explored: i) the extent to which covariance patterns change during postnatal ontogeny; ii) in which ontogenetic stages covariance patterns differ the most, and iii) the extent to which the phenotypic covariance pattern at different ontogenetic stages can be explained by the same processes determining additive genetic covariance. We sampled postnatal ontogenetic series for both marsupials, and placentals. Within each ontogenetic series, we compared covariance matrices (**P**-matrices) at different ontogenetic stages. Furthermore, we compared these **P**-matrices to two target matrices [adult **P**-matrix and an additive genetic covariance matrix (**G**-matrix)]. Our results show that for all ontogenetic series, covariance patterns from weaning onward are conserved and probably shaped by the same processes determining the **G**-matrix. We conclude that irrespective of eventual differences in how selection operates during most of postnatal ontogeny, the net response to such pressures will probably not be affected by ontogenetic differences in the covariance pattern.

## Introduction

Ontogenetic development is a multi-layered phenomenon in which developmental processes on a given stage act on the substrate laid out by processes that preceded them (Hallgrímsson et al., 2009). This sequential overlap leads to changes in the amount and distribution of morphological variation through time (Mitteroecker and Bookstein, 2009; Zelditch et al., 2006). Since natural selection is contingent on the availability and organization of morphological variation (Lande, 1979; Lande and Arnold, 1983), changes in this variation between life history stages can affect how selection operates, and how populations respond to selection in different life stages (Wasserman et al., 2021). Therefore, a broader comprehension of evolution involves understanding to what extent morphological variation changes during ontogeny.

For example, consider a scenario in which a pair of traits are associated (i.e. high integration *sensu* Olson and Miller, 1958) in the juvenile phase, but in the adult phase these traits are much less integrated (Figure 1; e.g., Sydney et al. (2012)). If selection operates on a single trait at the juvenile stage, evolutionary responses will be aligned with the major direction of variation of juveniles, leading to a correlated response in the second trait, even in the absence of trait association in the adult phase. Furthermore, in this scenario, the reconstruction of selection using the adult stage would suggest that selection is acting on multiple traits simultaneously, while in fact it is acting on a single trait earlier in development. Conversely, if the covariance patterns are relatively stable throughout ontogeny, selection would produce evolutionary responses that are similar across ontogenetic stages. Therefore, understanding how the variance is distributed on different ontogenetic stages can provide further insight about how complex phenotypes might evolve in response to natural selection.

**Figure 1:**
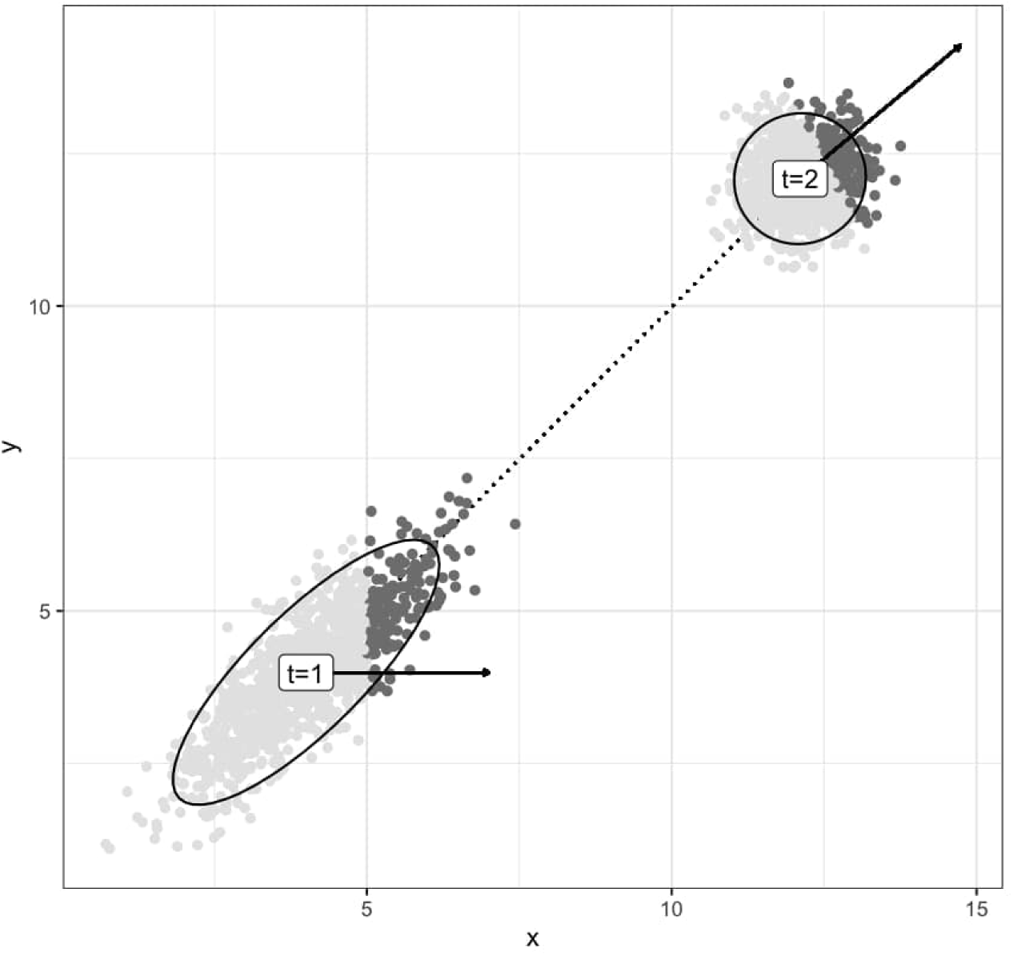
Effect of ontogenetic changes in covariance patterns on the evolutionary response of life-stage specific selection. Population is sampled in two moments: t=1 with strong integration and t=2 with weak integration. At t=1, the selection gradient (solid arrow) affect only the trait x (dark gray specimens), resulting in a selection differential that is correlated between x and y due to the high integration. At t=2, because x and y are not correlated, the reconstructed selection gradient (solid arrow) indicates that both traits where co-selected (dark gray specimens), while in fact, only x was.

The mammalian skull is a common model system for investigating the evolution of complex structures (e.g., Goswami, 2006; Haber, 2014; Machado et al., 2018), and there is compelling evidence that skull covariance patterns change during ontogeny (Atchley, 1984; Coleman et al., 1994; Goswami et al., 2012; Hallgrímsson et al., 2009; Mitteroecker and Bookstein, 2009; Mitteroecker et al., 2012; Nonaka and Nakata, 1984; Sydney et al., 2012; Zelditch, 1988; Zelditch et al., 1992; Zelditch and Carmichael, 1989; Zelditch et al., 2006). Despite this, the majority of studies on the evolution of the mammalian skull morphology focus only on adult phenotypes. Because selection is the net result of pressures acting throughout life-history stages, ignoring these developmental changes in trait association might affect our understanding of the evolution of this structure.

Here, we explore whether the observed variation in covariance patterns during postnatal ontogeny can impact the response to selection in age-structured populations under directional selection by assessing: i) the extent to which covariance patterns change during postnatal ontogeny; ii) in which ontogenetic stages covariance patterns differ the most (if they do at all), and iii) the extent to which the phenotypic covariance pattern at different ontogenetic stages mirrors the additive genetic covariance matrix (**G**-matrix). We developed a cross-sectional study of cranial trait covariances based on a sample of mammals with different developmental strategies. We sampled the Didelphimorphia marsupials *Didelphis virginiana* and *Monodelphis domestica*, the precocial platyrrhine primate *Sapajus apella*, and the altricial sigmodontinae rodent *Calomys expulsus* in different age classes encompassing the first months of life after birth to adulthood. We quantified covariance patterns of cranial morphological traits, and used published estimates of additive genetic covariance matrices for the same species or closely related taxa. Then, we compared covariance patterns among age classes within each one of these ontogenetic series to evaluate if observed differences would impact differentially the evolutionary responses under selection.

## Methods

### Sample

Our sample is composed of 1883 specimens belonging to five ontogenetic series: *Didelphis virginiana* and *Monodelphis domestica* (Marsupialia, Mammalia), *Sapajus apella* (Primates, Placentalia, Mammalia), and *Calomys expulsus* (Rodentia, Placentalia, Mammalia; Figure 2). Studied specimens are deposited in the following institutions: American Museum of Natural History (New York, USA), Field Museum of Natural History (Chicago, USA), Museu Nacional (Rio de Janeiro, Brazil), Museu Paraense Emilio Goeldi (Belém, Brazil), Museu de Zoologia da Universidade de São Paulo (São Paulo, Brazil), Museum of Vertebrate Zoology (Berkeley, USA), National Museum of Natural History (Washington D.C, USA), and Texas Biomedical Research Institute (San Antonio, USA).

**Figure 2:**
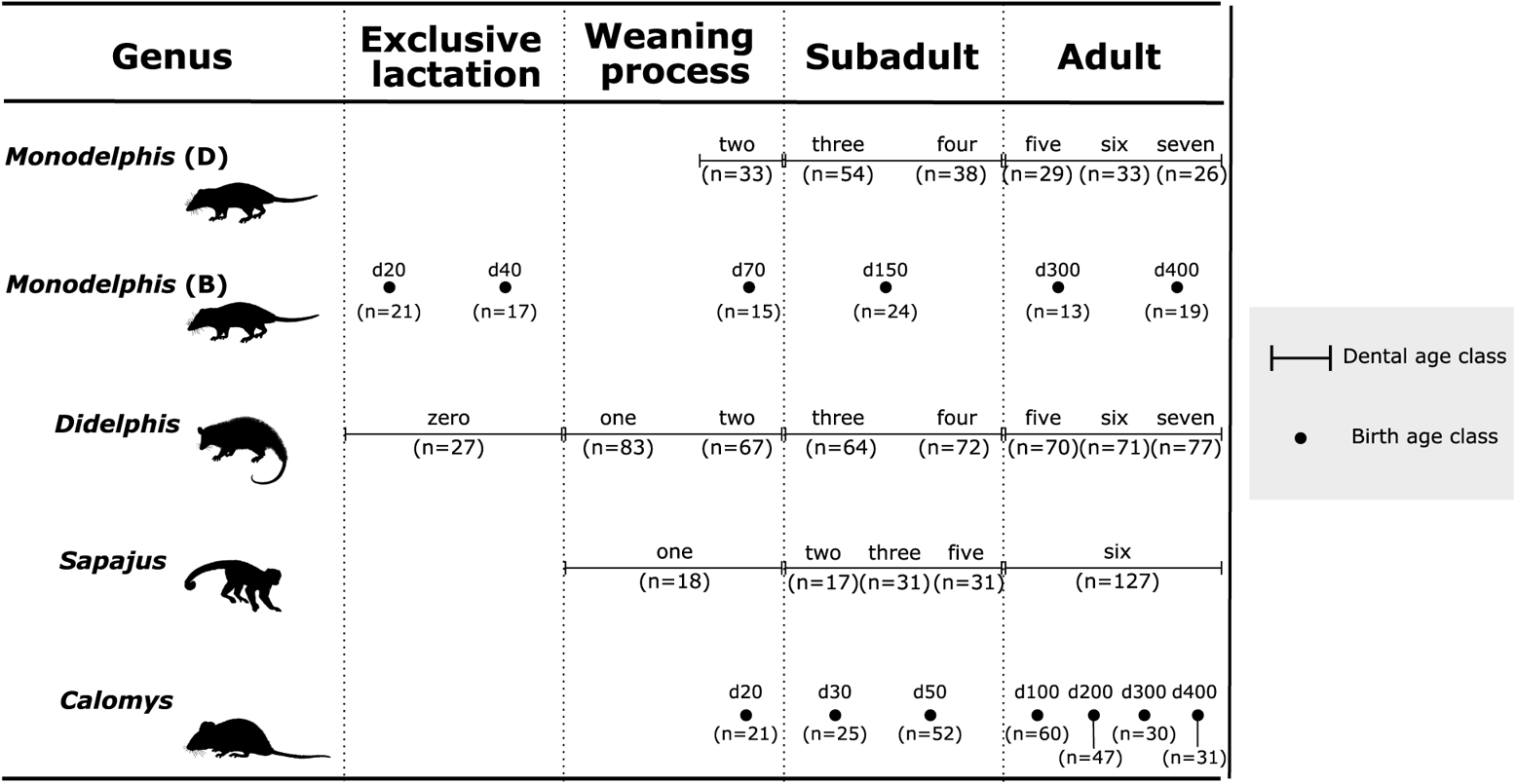
Schematic representation of the age classes and sample sizes (between parenthesis) for each ontogenetic series in relation to major life-history phases. Birth age classes are represented by dots and dental age classes by horizontal bars. The position of these symbols and the length of the bars are illustrative and are intended to show the broad distribution of data over the major life-history phases and differences in both sampling strategies (dental and birth age classes). Figures for each ontogenetic series are adapted from (Eisenberg, 1989; Eisenberg and Redford, 1999; Redford and Eisenberg, 1992). Figures not to scale.

We sampled specimens that were either wild caught (*Didelphis*, *Monodelphis*, and *Sapajus*), or derived from captive-bred colonies kept under stable controlled conditions [*Calomys* (Garcia et al., 2014), and *Monodelphis* (Porto et al., 2015)]. The two independent data sets for *Monodelphis* were labeled *Monodelphis* (D) for the wild-caught specimens and *Monodelphis* (B) for the captive-bred specimens. The acronyms stand for dental age class and birth age class, respectively, as explained below.

We classified our specimens according to age classes. Wild-caught specimens were classified according to dental eruption and wear (i.e., dental age class), while captive-bred specimens were classified according to days after birth (i.e., birth age class; Figure 2). Dental age class for *Didelphis* and *Monodelphis* (D) were determined based on maxillary dental eruption and wear (Tribe, 1990; Tyndale-biscoe and Mackenzie, 1976). We added an extra class (zero), composed of specimens with no erupted teeth. For *Didelphis*, the ontogenetic stages sampled most likely include lactation (dental age class zero), the start of solid food ingestion (dental age class one), end of the weaning (between dental age classes one and two), to adulthood [dental age class above four; Abdala et al. (2001); McManus (1974); Sebastião and Marroig (2013); Tyndale-biscoe and Mackenzie (1976); van Nievelt and Smith (2005)].

For *Monodelphis* (D), the ontogenetic stages sampled most likely include the end of the weaning process (during dental age class two) to adulthood [dental age class above four; Sebastião and Marroig (2013); van Nievelt and Smith (2005)]. *Monodelphis* (B) specimens were classified according to birth age classes 20, 40, 70, 150, 300, and 400 days after birth. These classes encompass exclusive lactation (20 and 40), to the very end of the weaning process (70), to adulthood [300 and 400; Nievelt and Smith (2005)]. The birth age classes are analogue to the following dental age classes: 20 = zero; 40 = zero to one; 70 = two; *≥* 150 = *≥* four (Nievelt and Smith, 2005; van Nievelt and Smith, 2005). The dental age classes for *Sapajus* were determined based on the premaxillary and maxillary dental eruption (Richtsmeier et al., 1993). They span from weaning (dental age class one) to adulthood [dental age class six; Fragazy et al. (2004); Marroig and Cheverud (2001)]. The *Calomys* were classified according to the birth age classes 20, 30, 50, 100, 200, 300 and 400 days after birth. Specimens comprise from around weaning (20 days) to adulthood [100 days onwards; Hingst-Zaher et al. (2000)].

For *Sapajus*, almost all specimens we studied are assigned to *S. apella* (n = 200), however 24 specimens are of uncertain classification (*Sapajus* sp.). These specimens belong to the age classes one (n = 6), two (n = 11) and three (n = 7). Inspections of Principal Component Analysis (PCA) plots showed that specimens with unknown species fell within the distribution of *S. apella*, cluster along other juveniles. (Figure A1). A non-parametric multivariate analyses of variance (NP-MANOVA, Anderson, 2001; Collyer and Adams, 2018; McArdle and Anderson, 2001) using size (to control for ontogenetic variation) and species assignment as factors showed no differentiation between *S. apella* and *Sapajus* sp. (Table A1). Based on these results, we pooled both groups in our analysis to improve sample sizes.

### Landmarks and Measurements

Traits on this study are linear distances derived from the 3D coordinates of 32 homologous landmarks measured on all specimens using a 3D digitizer. All specimens were measured with the same instrument (Microscribe MX; Immersion Corporation, San Jose, California) with the exception of adult *Sapajus* that were measured with a 3Draw digitizer (Polhemus Inc., Colchester, Vermont). Based on these landmarks, 35 linear distances were calculated in millimeters [for details related to the landmarks and distances acquisition refer to (Cheverud, 1995; Porto et al., 2009; Shirai and Marroig, 2010)]. The set of distances calculated aimed to represent the whole cranium morphology and important developmental and functional relationships among cranial regions, while avoiding redundancy (Cheverud, 1982; Marroig and Cheverud, 2001).

Some sources of measurement error could introduce non-biological variance in our data sets. First, more than one equipment was used to collect data. However, tests performed with adult specimens measured with both devices indicated that this source of variation is negligible (G. Marroig, *pers. comm.*). Second, specimens were measured by different observers. *Didelphis* and *Monodelphis* (D) crania were measured by HS. *Monodelphis* (B) crania were measured by AP. *Calomys* crania were measured by GG, and *Sapajus* crania were measured by GM. Nevertheless, all specimens were measured following the same protocol, and since we are studying covariances within each ontogenetic series, and all specimens per ontogenetic series were measured by the same person, we expect inter-observer error to be irrelevant.

Lastly, to evaluate the possible effect of within-sample measurement error in covariance estimates we calculated trait repeatabilities (Lessells and Boag, 1987) for samples which were measured twice, namely *Didelphis*, *Monodelphis* (D), and *Calomys*. Measurement errors were calculated for each ontogenetic series at each age class with more than 14 specimens for each linear distance independently. In most cases, measurement errors are negligible, since most repeatabilities were high (*>* 0.8; Figure A2). Low repeatabilities (*<* 0.8) were observed for traits exhibiting low variances (Figure A3). These traits were also very short, approaching the spatial resolution of the digitizer (Figure A3). In all subsequent analyses, specimens’ traits were represented by the mean of replicated measurements. Although the *Sapajus* and the *Monodelphis* (B) specimens were not measured twice, the measurement errors observed for other Platyrrhini measured by GM (Marroig and Cheverud, 2001), and for smaller Didelphimorphia measured by AP (Porto et al., 2009) were negligible. Therefore, we assume the *Sapajus* and *Monodelphis* (B) specimens also presented negligible measurement errors.

### Phenotypic covariance matrices

Our study is concerned with how the association among traits changes during ontogeny. To quantify trait associations, we calculated phenotypic covariance matrices (**P**-matrices) of each age class per ontogenetic series. Because our sample includes both male and female specimens (Table A2), we evaluated if the presence of sexual dimorphism could affect our covariance estimation. To verify this possibility, we used the following approaches. First, we used pairwise NP-MANOVA (significance at *p*(*α*) *<* 0.05) to evaluate the effect of sex on morphology for each age class. In cases of insufficient sample sizes for NP-MANOVA we used pairwise non-parametric univariate analysis of variance (NP-ANOVA), and considered sexual dimorphism to be present whenever two or more traits had significantly covaried with sex at *p*(*α*) *<* 0.01. Second, sexual dimorphism was also graphically evaluated using Principal Component Analyses. Lastly, we assessed the impact that controlling for sexual dimorphism have on the covariance estimates by evaluating the extent to which the covariance structure is altered by the exclusion of sex in the analyses. To do that, first we calculated matrices with and without controlling for sexual dimorphism and compared them using the Random Skewers method (for details on the method see below) and also calculating the difference between the trace of the matrices. Because sexual dimorphism was identified as a source of variance in at least one age class per ontogenetic series, we calculated the residual pooled within-group **P**-matrices for all samples using the general linear model approach (Marroig and Cheverud, 2004). This step is important because if an effect is a strong source of variance, it distorts matrix estimates, which will reflect inter-group (e.g., female and male differences) instead of intra-group covariances [see Figure SI1 in Machado et al. (2018)]. For the adult age class, we also controlled for the effect of the age classes involved (Figure 2), as described above.

Any matrix is estimated with error, and the lower the ratio between the number of trait and sample size the worst is the estimation. We quantified error in matrix estimation by using a Bayesian posterior sample of the covariance matrices using a multivariate normal likelihood function on the residuals of the linear models and an inverse Wishart prior on the covariance structure. The prior covariance matrix is set to a diagonal matrix with the observed variances on the diagonal, and the prior degrees of freedom is set to the number of traits (*k* = 35). This particular choice for the likelihood and prior distribution can be solved analytically and results in a posterior distribution from which we can sample directly (Murphy, 2012). Furthermore, this method has the added benefit of ensuring that the matrices from the posterior sample are positive-definite and, therefore, invertible. We took 100 posterior samples for each age-class from each ontogenetic series, producing 3,600 matrices in total.

To visualize differences and similarities between the **P**-matrices we performed a Principal Coordinate Analysis (PCoA). The PCoA generates a lower-dimensional representation of a multivariate dataset similarly to the Principal Component Analysis (PCA). Different from PCA, the PCoA is based on the spectral decomposition of the double-centered distance matrix that represents the dissimilarities among samples. The eigenvectors of this analysis (Principal Coordinates; PCo) express the scores of each sample on this reduced space, with the leading eigenvectors representing the axes in which covariance matrices differ the most, and the latter representing the axes in which they differ the least. The dissimilarity between matrix was calculated based on the Riemmanian distance, which is the metric of the space of square symmetric positive definite matrices (Bookstein and Mitteroecker, 2014; Le Maître and Mitteroecker, 2019; Mitteroecker, 2009). Because Matrices Riemmanian distances are sensitive to scale, matrices were set to have the same size (trace=1) prior to the calculation. The resulting PCo space then only relates to matrix shape.

### Additive genetic covariance matrices

The **P**-matrix is determined by the additive genetic covariance matrix (**G**-matrix), plus the environmental covariance (Falconer and MacKay, 1996). The **G**-matrix quantifies the genetic contribution to trait’s patterns of inheritance and co-inheritance (covariance), and is essential for predicting multivariate evolution of these traits (Falconer and MacKay, 1996; Lande, 1979; McGuigan, 2006). Because of that, we evaluated if patterns of covariance quantified in our age-specific **P**-matrices can be explained by the distribution of heritable variation encoded in the **G**-matrix. To do this, we compared our age-specific **P**-matrices within ontogenetic series with estimated target **G**-matrices (see details regarding the comparison method below). The reasoning behind this approach is that since **G**-matrices of complex traits represent the net-effect of multiple pleiotropic effects channeled through developmental pathways (Cheverud, 1996a), finding a high similarity between the age-specific **P**-matrices and the target **G**-matrix means that trait associations within **P**-matrices are probably determined by the same processes determining the **G**-matrix pattern.

Depending on the ontogenetic series, a different target **G**-matrix was adopted. For comparisons within Didelphimorphia [*Didelphis*, *Monodelphis* (D), and *Monodelphis* (B)] we used a matrix for *Monodelphis domestica* (Porto et al., 2015) and for the comparison within *Calomys*, we used a matrix estimated for the same species (Garcia et al., 2014). These matrices were estimated using the same individuals from the *Monodelphis* (B) and *Calomys* ontogenetic series, respectively. There is no available **G**-matrix for *Sapajus*. Therefore, we used a matrix estimated for *Saguinus* (Cheverud, 1996b). The *Saguinus* **G**-matrix is highly similar to **P**-matrices for adult samples of all New World Monkey genera, including *Sapajus* (Marroig and Cheverud, 2010). Thus, this **G**-matrix can be considered a good rough approximation of the **G**-matrix for *Sapajus*, at least for patterns of covariance since differences due to scale (*Saguinus*: 400 g; *Sapajus*: 2800 g) result in larger variances and covariances for *Sapajus*.

The **G**-matrix for *Calomys* was estimated from 365 specimens comprising individuals of both sexes and of different age classes, and that were raised in an unbalanced colony design (i.e., containing both paternal and maternal half-sibs). The **G**-matrix for *Monodelphis* was estimated from 199 adult specimens belonging to 16 partially inbred strains. Both matrices were estimated using a Bayesian sparse factor model (Runcie and Mukherjee, 2013), in a full animal model for *C. expulsus* and a structured random effect model for *M. domestica* that used the genetic distance between strains to define the covariance between random effect levels. For *Saguinus*, the **G**-matrix was estimated from 462 specimens pooled from two different species: *S. oedipus* and *S. fuscicollis*.

Colony designs were unbalanced, including full and half-siblings, and various kinds of collateral relatives. The matrix was estimated under a Maximum Likelihood framework in a full animal model (Cheverud, 1996b; Konigsberg and Cheverud, 1992). In all cases, unwanted sources of variation, like sex, were controlled using fixed effects.

### Matrix comparisons

To evaluate how much covariance patterns change during ontogeny, we compared age-specific **P**-matrices within ontogenetic series using two different approaches: the Krzanowski Subspace Comparison for multiple matrices [KC; (Aguirre et al., 2014; Krzanowski, 1979) and the Random Skewers [RS; (Cheverud and Marroig, 2007)].

We used KC to compare age-specific **P**-matrices within each of the five ontogenetic series (Figure 2). KC is a global test of similarity among all matrices, and measures the alignment of the morphospaces spanned by the first few eigenvectors of the matrices being compared. Structurally similar matrices should have most of their variation in a similar subspace. The matrix that describes the common subspace is defined as

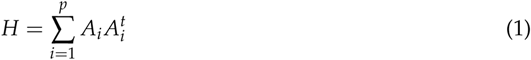

where *A_i_* is a column matrix containing the first *k* = *n*/2 *−* 1 eigenvectors of the *i*-th matrix being compared, *p* is the number of matrices being compared, and *^t^* denotes matrix transposition. The eigenvalues of *H* are at most *p*, and any eigenvector of *H* whose associated eigenvalue is equal to *p* can be reconstructed by a linear combination of the eigenvalues included in the *A_i_* matrices, and so is shared by all the matrices.

To create a null distribution for the eigenvalues of *H*, we used a permutation approach (Aguirre et al., 2014), randomizing the residuals from the fixed effect models used to calculate the P-matrices between the age classes within lineage 1000 times. For each permuted sample, we repeat the Bayesian posterior sampling of the P-matrix in each age class and calculate the *H* matrix. This provides a null distribution for the eigenvalues of *H* under the hypothesis that the residuals for each age class came from the same population. The confidence intervals in the observed *H* matrix are obtained using the posterior distribution of the true P-matrices, while the permuted confidence interval under the null hypothesis combines the uncertainty from the randomization and from the posterior distributions for each randomized sample. If the observed eigenvalues of *H* were significantly different from the randomized eigenvalues, we concluded that the matrices from each age class have a different structure, otherwise, we concluded that the matrices are similar.

Additionally, we used the RS method to compare the median of each posterior distribution of age-specific **P**-matrices within each ontogenetic series against two target matrices: the median of the posterior distribution of the adult pooled within-group **P**-matrix of each ontogenetic series and a **G**-matrix (see above for details on the **G**-matrices compared in each case). RS is based on the following equation:

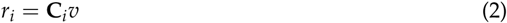

where *v* is a random vector, **C_i_** is a matrix being compared and *r_i_* is the response vector. The RS similarity is then defined as the mean vector correlation between the response vectors obtained by applying the same set of random vectors to two covariance matrices (Cheverud and Marroig, 2007; Melo et al., 2015). If the covariance matrices have similar structures, their response vectors will be closely aligned and the RS will be close to 1. If they have unrelated structures, the direction of the response will be different and the RS will be close to zero.

For reasonably similar matrices, as in our case, *r* is strongly and negatively correlated to the Riemannian distance (Figure A4), with the advantage that, under certain conditions, RS has a straightforward biological interpretation. The RS equation has the same format as the multivariate response to selection equation Δ*z* = **G***β* (Lande, 1979), where *β* is the selection gradient (direction of maximum fitness increase), **G** is the **G**-matrix and Δ*z* is the evolutionary response to selection vector. Thus, for evolutionary studies, RS is a measure of the average alignment between the evolutionary responses of two populations subjected to the same selective pressures. We can calculate this if we have access to the **G**-matrices or to **P**-matrices that are good proxies for the corresponding **G**-matrices. This is why we compared the age-specific **P**-matrices within ontogenetic series not only with adult **P**-matrices, which allows us to scrutinize phenotypic differences in covariance patterns within the ontogenetic series, but also with **G**-matrices.

As explained above, if the RS for age-specific **P**-matrices within ontogenetic series are similar to the target **G**-matrices, we can infer that the phenotypic covariance patterns within each ontogenetic series can be sufficiently explained by the same processes determining the (co)inheritance of traits. Furthermore, this similarity would also suggest that different age classes respond similarly to selection. The proportionality between **P**- and **G**-matrices, sometimes referred to as the *Cheverud Conjecture* (Roff, 1995), has been verified for adult skull traits in many lineages of mammals (Cheverud, 1995; Hubbe et al., 2016; Machado et al., 2018; Marroig and Cheverud, 2001; Oliveira et al., 2009; Porto, 2009; Shirai and Marroig, 2010), and specifically for the lineages investigated here (Cheverud, 1995; Garcia et al., 2014; Marroig and Cheverud, 2010; Porto et al., 2015). Thus, by evaluating the RS similarity between age-specific **P**-matrices within ontogenetic series and target **G**-matrices, we are also testing if the Cheverud Conjecture can be extended to other ontogenetic stages besides the adult one.

Due to sampling error associated with the estimation of covariance matrices, the maximum RS value (*r_max_*) between two matrices will never be one, even if the underlying samples come from the same population. To account for this, we calculated matrices repeatabilities (*t*), which is a measure of the expected similarity between the true underlying covariance structure and the one calculated from the sample. The *t* value for all matrices was determined using a Monte Carlo resampling procedure of self-correlation (Marroig and Cheverud, 2001; Porto et al., 2009). For every covariance matrix, 1,000 Monte Carlo samples were made keeping sample size constant. Repeatabilities for the *Calomys* and *Monodelphis* **G**-matrices were calculated using the published effective sample sizes, and the one for *Saguinus* was taken from the literature (Cheverud, 1996b; Marroig and Cheverud, 2010). Covariance matrices were estimated for each of the resamples and RS was used to compare the original and the resampled matrices. The *t* value was then obtained as the mean RS value between original and resampled matrices. Adjusted RS similarity was then estimated as *r_adj_* = *r_obs_*/*r_max_*, where *r_obs_* is the observed similarity among samples and *r_max_* is the geometric mean of *t*s of the pair of matrices being compared (Cheverud, 1996b). The procedure described above provides inflated estimates of repeatabilities for poorly estimated **P**-matrices, thus providing conservative corrections for RS when at least one poorly estimated **P**-matrix is considered in the analysis.

Lastly, it is important to notice that both KC and RS are primarily concerned with the directions of variance in the morphospace. Magnitudes of variance are strongly correlated with the scale of the organisms, which changes considerably across different age classes. More specifically, KC considers only the eigenvectors (directions) of the matrices under study, disregarding the corresponding eigenvalues (variance in these directions); and RS considers the direction and magnitude of covariance patterns, but quantifies only the alignment (directions) of response vectors, and not their length (magnitude). In other words, the RS similarity depends on the relative distribution of variances, not on its magnitude.

### Statistical analyses

All analyses were done in the R Core Team (2019) programming environment. The EvolQG v0.3-1 (Melo et al., 2015) package was used for matrix estimation, RS and KC comparisons, the package RRPP (Collyer and Adams, 2018, 2019) was used for the NP-ANOVAs and MANOVAs and the package vcvComp (Le Maître and Mitteroecker, 2019) was used for the PCoA. Analyses were done independently by three authors (FAM, DM and GG), and results were consistent between runs.

## Results

The first two leading eigenvectors of the PCoA explained 29.97% and 4.11% of the total variation in the sample, respectively (Figure 3). PCoA1 separates *Didelphis* and *Sapajus* samples with higher scores from *Monodelphis* (B), *Monodelphis* (D) and *Calomys* samples with lower scores. PCoA2 shows a contrast between *Monodelphis* (B), *Monodelphis* (D), and *Didelphis* with lower scores from *Calomys* and *Sapajus* with higher ones. PCoA1 presents some ontogenetic structuring of the marsupial species, with younger age classes showing lower values than adult classes. Furthermore, the *Monodelphis* (B) and *Monodelphis* (D) form almost a continuum, with latter stages of *Monodelphis* (B) neighboring intermediary to late stages of *Monodelphis* (D). The following eigenvectors explained *<* 3% of the total variation and show no clear taxonomic or ontogenetic structure (Figure A5).

**Figure 3:**
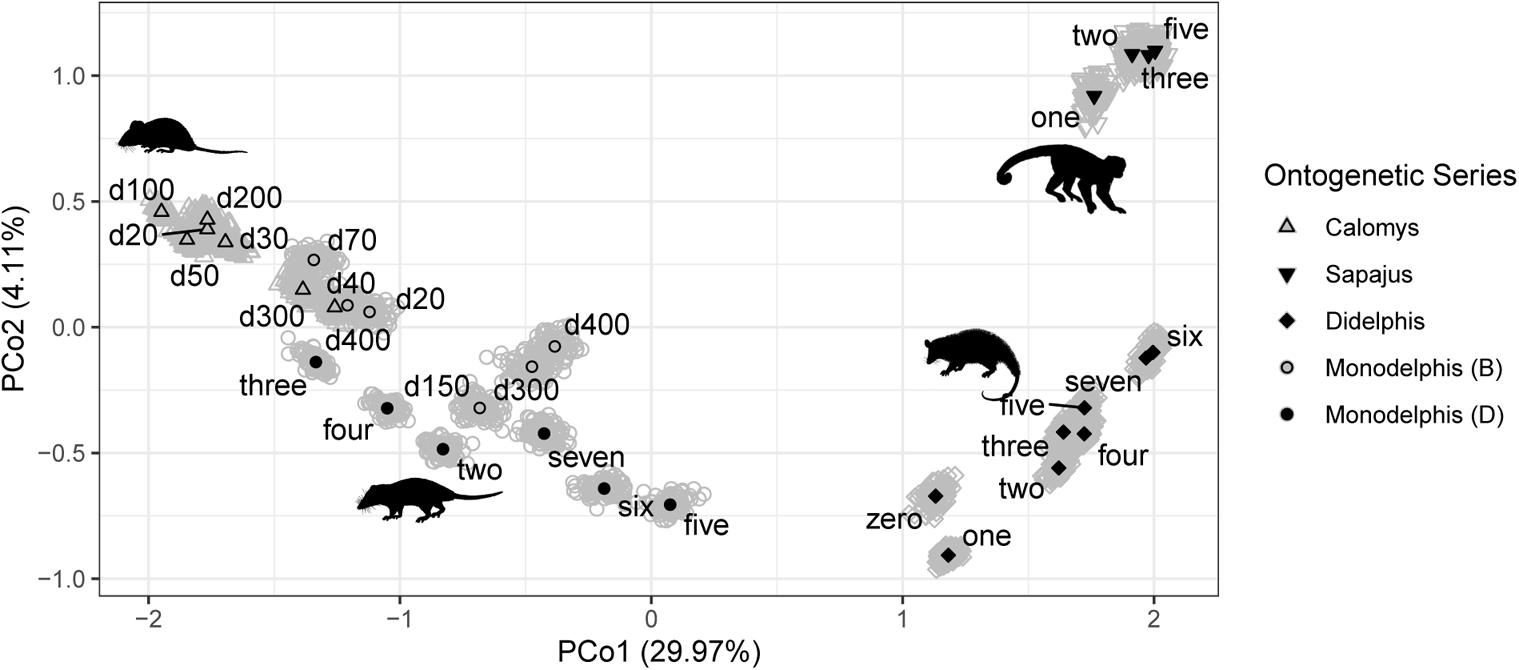
Distribution of age-specific **P**-matrices on the first two leading principal coordinates based on the Riemannian distance. Black symbols represent the median of each posterior distribution of age-specific **P**-matrices within each ontogenetic series. Gray symbols represent 100 matrices from the posterior distribution of age-specific **P**-matrices within each ontogenetic series.

The KC analysis shows that the posterior eigenvalue distribution fully overlaps with the null-distribution for all ontogenetic series (Figure 4), suggesting that, despite the dispersion observed in Figure 3 for some age classes, covariance structures tend to be similar with ontogenetic series. The only possible exception is within the *Monodelphis* (B), where the posterior distribution of the leading eigenvalue falls partly outside of the null-distribution (Figure 4B).

**Figure 4:**
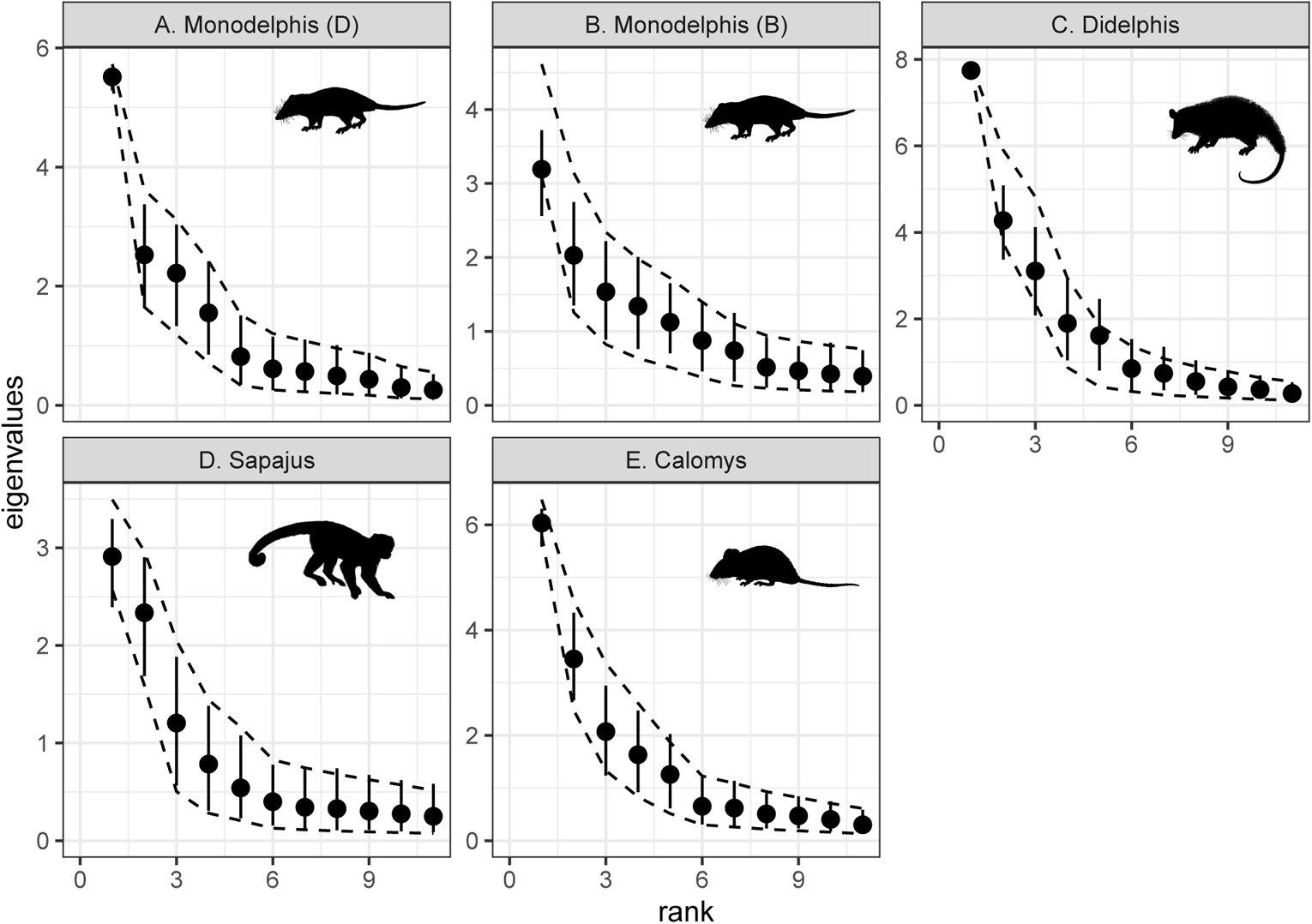
Distribution of the empirical (bars) and null (dashed lines) distributions of the eigen-values of the Krzanowski Subspace Comparison for each ontogenetic series. Overlap between empirical and null eigenvalue distributions indicates that observed covariance matrices are as similar as matrices in which age-class was randomized across individuals. While not identical, this is evidence that observed matrices are compatible with a single underlying covariance structure across age classes.

The pairwise RS analysis is consistent with this interpretation for most of the sample (Figure 5), showing that comparisons between age-specific **P**-matrices and the adult **P**-matrix yielded high similarity values (*RS >* 0.81), the exceptions being the age classes d20 and d40 for *Monodelphis* (B) (*RS* = 0.64, *RS* = 0.73, respectively). The RS comparison between age-specific **P**-matrices and target **G**-matrices presented a very similar result to the comparison with adult **P**-matrices, except that similarity values tended to be slightly lower (*RS >* 0.69). Exceptions were *Sapajus*, which wielded consistently lower similarity values than the comparison with the adult’s **P**-matrix (0.68 *< RS <* 0.79), and *Monodelphis* (B), which showed lower values in general, and particularly for the two younger age classes (*RS <* 0.44). In the case of *Sapajus*, these lower similarities are still remarkably high considering that they are being compared to a **G**-matrix estimated for a species of a different family (Callitrichidae). With the exception of *Monodelphis* (B), there is a trend of comparisons involving poorly estimated matrices presenting lower RS values, which is expected given the contribution of noise to the covariance patterns added by sampling error. Nevertheless, even the comparisons involving these matrices are fairly high.

**Figure 5:**
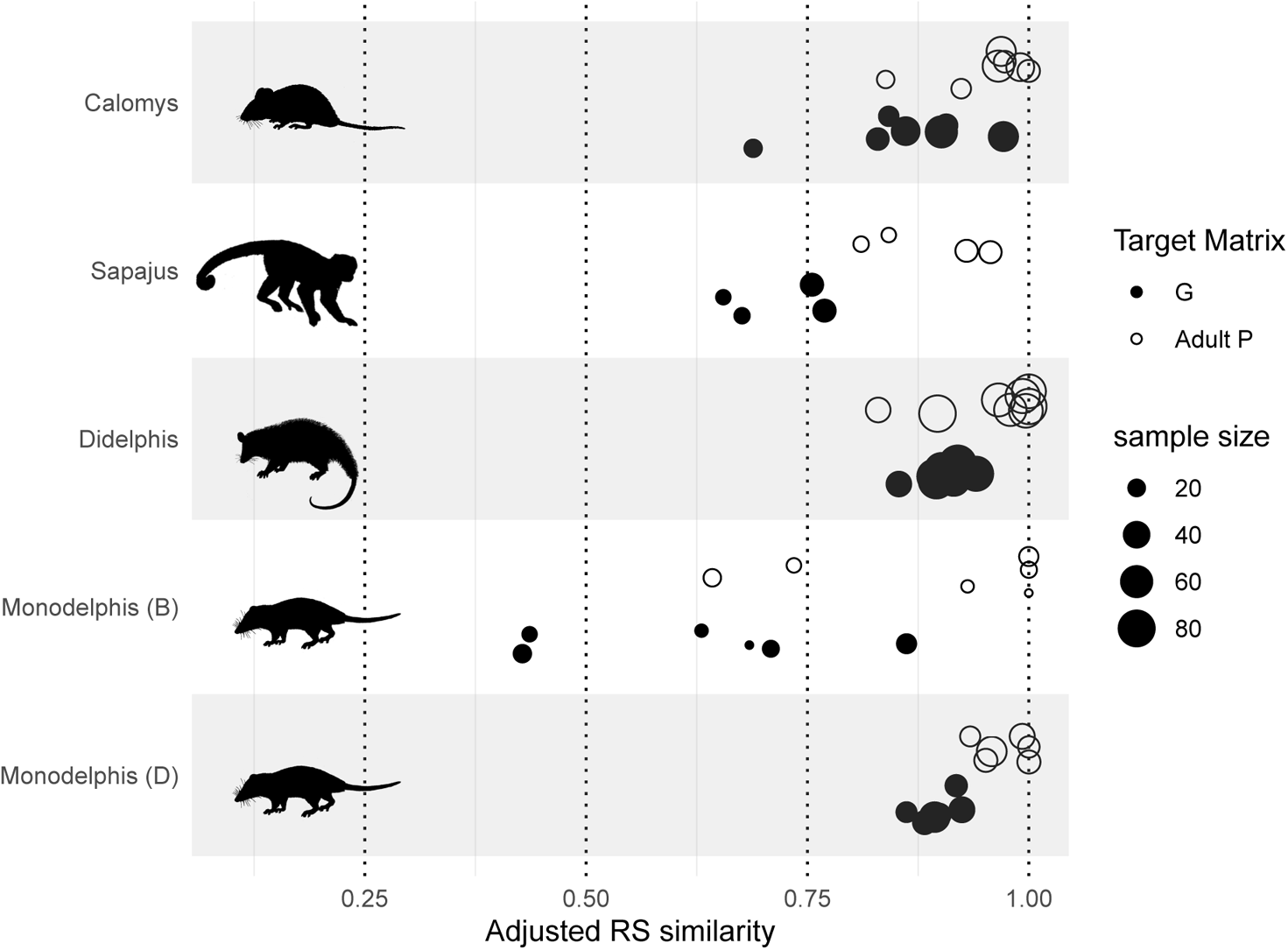
Distribution of adjusted Random Skewers similarity between age-specific covariance matrices and adult phenotypic matrix or a target additive genetic covariance matrix.

## Discussion

In this contribution, we have investigated how the covariance pattern of mammalian skull traits changes during the postnatal ontogeny in five ontogenetic series of four mammalian species, which show substantial life-history and pre and postnatal development differences (Smith, 1997). Covariance patterns from weaning onward are fairly stable within ontogenetic series (Figure 4, 5), and these patterns are largely driven by the same processes governing the (co)inheritance of traits (Figure 5), which suggest that the Cheverud Conjecture holds for covariance patterns during all post-weaning development, not only for adults as previously reported (Akesson et al., 2007; Cheverud, 1988, 1996b; House and Simmons, 2005; Porto et al., 2009; Reusch and Blanckenhorn, 1998; Roff, 1995). Furthermore, the extension of the Cheverud Conjecture throughout the ontogeny suggests that similar selective pressures operating on different life history stages will probably result in a similar evolutionary response.

One potential exception to this pattern is *Monodelphis* (B). While the KC failed to find any significant difference between age classes, this ontogenetic series was the only case where a substantial part of the posterior distribution of eigenvalues of the common subspace matrix (**H**-matrix) fell outside the null distribution (Figure 4B, first eigenvalue). Furthermore, the leading axis of the PCoA is mainly associated with ontogenetic differences within *Monodelphis*, and Didelphimorphia to a lesser degree (Figure 3) and the RS showed differences during the lactation phase, at birth age classes d20 and d40 (Figure 5). At these early postnatal stages, covariance patterns were shown to be different to some extent from the patterns of additive genetic covariance as well (Figure 5). This suggests that at least for this group, the skull trait covariance is subject to changes during early stages of postnatal development, which stabilizes only around weaning. Even though our sample sizes at this early postnatal stage are among the lowest in our sample, making their matrices the worse estimates we have, these results seem to be at least in part real biological signals (Figure A6). Nevertheless, probably due to low sample sizes, specific investigation into how trait covariances are changing between age-classes are unfortunately inconclusive (Figure A7,A8).

In contrast, *Didelphis* during lactation (dental age class zero) presented similar covariance patterns with older dental age classes, but this result could be the consequence of sampling specimens based on dental age classes (Figure A9). Specifically, by pooling individuals with different absolute ages within the same class, one might be artificially inflating the effect of size, forcing eigenvectors to align themselves with those of latter stages, which are usually dominated by size variation. Consequently, in such cases, RS will detect high similarities between matrices (Porto et al., 2013; Rohlf, 2017). In fact, the leading eigenvector of the age class zero for *Didelphis* shows almost all loadings with the same signal, as expected for size (Jolicoeur, 1963), while the same is not true for age classes d20 and d40 of *Monodelphis* (B) (Figure A10). For placental mammals (*Sapajus* and *Calomys*), we do not have samples before weaning (Figure 2), so it is unclear if the same pattern observed on *Monodelphis* could be extended to all mammals. More studies focusing on earlier ontogenetic stages, and on larger samples based on birth age classes will be required to better understand this phenomenon.

At this point, we cannot further discuss potential changes in covariance patterns prior to weaning, and we can only speculate on the reasons for the maintenance of covariance pattern from weaning onward. The mammalian postnatal ontogeny occurs over major life-history phases (Abdala et al., 2001; McManus, 1974; Nievelt and Smith, 2005) that may influence development (Atchley, 1984; Hallgrímsson et al., 2009; Sibly et al., 2014; Zelditch, 1988), such as lactation, beginning of solid food ingestion, and weaning. The postnatal stage is also characterized by several changes in the skull (Abdala et al., 2001; Flores et al., 2006; Zelditch et al., 1992), such as the faster growth of the viscerocranium in relation to the neurocranium, and the development of muscles of the masticatory apparatus to deal with solid food, which influences the growth of underlying bones (Kiliaridis, 1995). Nevertheless, most postnatal development is shaped by growth and muscle–bone interactions, which are relatively late developmental inputs (Hallgrímsson et al., 2009, 2007). Given that the covariance pattern observed in adult populations are the end result of several hierarchical developmental processes (Hallgrímsson et al., 2009, 2007), it is possible that events that occur later in development might have smaller effects on determining covariance patterns due to constrains imposed by early processes (Atchley, 1984) and thus may have limited influence over covariance patterns. Alternatively, these later stages can be shaping the covariance pattern, overwriting early developmental inputs (Hallgrímsson et al., 2009).

Irrespective of the reasons for the maintenance of covariance patterns from weaning onward, our findings have important implications for evolutionary, genetic, and ecological studies. We showed that during lactation covariance patterns may vary considerably, at least in marsupials, but that from weaning onward, covariance patterns become relatively stable. Thus, for specimens spanning from weaning to adulthood, selection operating on different postnatal ontogenetic stages might have similar consequences on the responses produced in terms of the pattern of changes in traits averages.

Given that during lactation individuals should be subjected to very different selective regimes than after weaning, our results are reassuring in that working with a single ontogenetic stage will not lead to misleading conclusions. We have sampled species that are distantly related and that show profoundly different pre- and postnatal developmental strategies. While future studies will no doubt be important to reevaluate this conclusion, we suggest that the broad taxonomic and life history range encompassed by our sample implies that our results can be extended to all therian mammals. Given that adult covariance matrices tend to be very similar among closely related groups (Hubbe et al., 2016; Machado et al., 2018; Marroig and Cheverud, 2001; Rossoni et al., 2019; Shirai and Marroig, 2010), there is little reason to believe that evolutionary lineages with more similar developmental patterns, such as within placentals or marsupials, will have a higher influence of developmental changes over covariance patterns.

## Conclusion

Our findings suggest that for mammals the study of the life-history changes and evolutionary consequences under selection (or genetic drift) is much facilitated by shared and common co-variance patterns among traits during most of postnatal ontogeny. Thus, even though selection might be operating in different directions during most of postnatal ontogeny, due to differences in life-history phases and fitness components, the net response to such pressures will probably not be biased by differences in the covariance pattern during the postnatal ontogeny.

While our findings support using a single weaning-onward ontogenetic stage in the investigation of selective pressures and evolutionary responses, they also highlight the need for a more comprehensive understanding of how covariances changes between birth and weaning. In addition, it is important to better understand species life histories to evaluate when and how selection is operating, selective explanations inferred from adult morphologies might be a consequence of selection operating on other life stages. This is particularly relevant if species show relatively drastic change in some ecological aspect during ontogeny [e.g., Drago et al. (2009); Tanner et al. (2010)]. Lastly, a full account of how changes in selective pressures during ontogeny can impact the response to selection would require considering the covariances for the same trait across ontogenetic stages as well. This requires longitudinal morphological data on the animals, which is a challenging but interesting venue for future research.

## Acknowledgment

The authors are thankful to people and institutions that provided generous help and access to collections: A. Fleming, R. MacPhee, R. Voss, and E. Westwig (American Museum of Natural History); B. Patterson, L. Heaney, and B. Stanley (Field Museum of Natural History); S. Costa, and J. de Queiros (Museu Paraense Emilio Goeldi); F. Barbosa, S. Franco, J. A. de Oliveira, and L. Salles (Museu Nacional); C. Conroy, E. Lacey and J. Patton (Museum of Vertebrate Zoology); M. Brett-Surman, R. Chapman, L. Gordon, and D. Lunde (National Museum of Natural History); J. Gualda, and M. de Vivo (Museu de Zoologia da Universidade de São Paulo); and J. VandeBerg (Texas Biomedical Research Institute). We are deeply indebted to two anonymous reviewers and editors Erol Akcay and Tim Connallon for their insightful comments on a previous draft of this manuscript. This research was supported by grants and fellowships from Fundação de Amparo à Pesquisa do Estado de São Paulo (FAPESP), Coordenadoria de Aperfeiçoamento de Pessoal de Nível Superior (CAPES), Conselho Nacional de Desenvolvimento Científico e Tecnológico (CNPq), Museum of Vertebrate Zoology (UC-Berkeley), American Museum of Natural History, Field Museum of Natural History, and NSF (DEB 1942717).

## Appendix A: Supplementary Tables and Figures

**Table A1:**
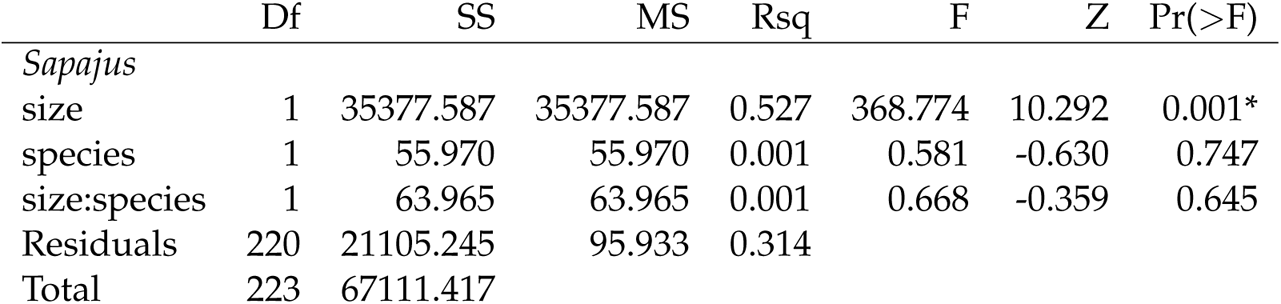
Non-parametric RRPP MANOVA results for the effect of size (measured as the geometric mean of traits) and species identity on trait variation for the *Sapajus* sample. Df-degrees of freedom. SS-Sum of squares. MS-Mean squares. Rsq-coefficient of determination. F-F statistic. Z-z-transformed effect sizes. Pr(*>*F-P-value based on 1000 permutations. *-effects considered significant at *α* = 0.05.

|  | Df | SS | MS | Rsq | F | Z | Pr(>F) |
| --- | --- | --- | --- | --- | --- | --- | --- |
| *Sapajus* |  |  |  |  |  |  |  |
| size | 1 | 35377.587 | 35377.587 | 0.527 | 368.774 | 10.292 | 0.001* |
| species | 1 | 55.970 | 55.970 | 0.001 | 0.581 | -0.630 | 0.747 |
| size:species | 1 | 63.965 | 63.965 | 0.001 | 0.668 | -0.359 | 0.645 |
| Residuals | 220 | 21105.245 | 95.933 | 0.314 |  |  |  |
| Total | 223 | 67111.417 |  |  |  |  |  |

**Table A2:**
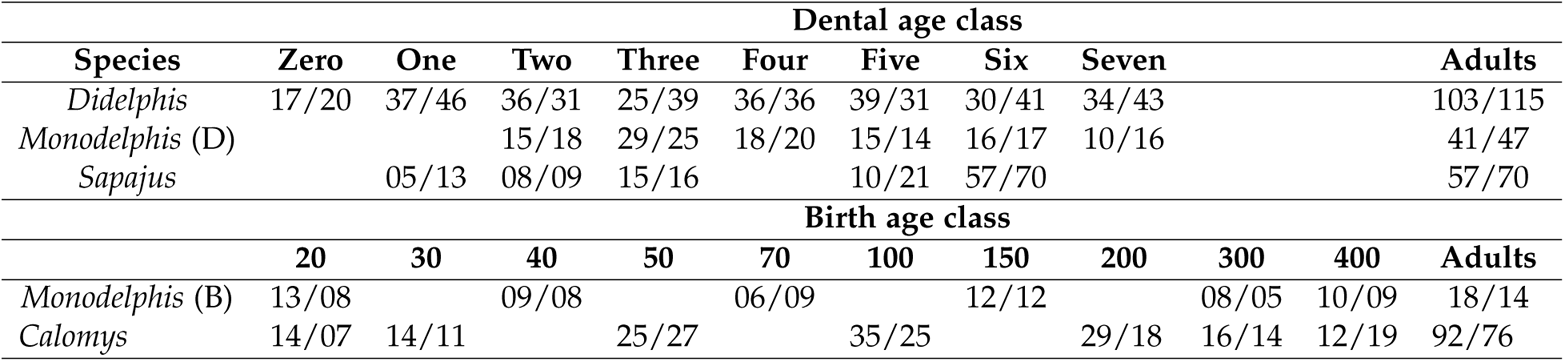
Sample sizes (Females/Males) for each age class and ontogenetic series.

| Dental age class |  |  |  |  |  |  |  |  |  |  |  |
| --- | --- | --- | --- | --- | --- | --- | --- | --- | --- | --- | --- |
| Species | Zero | One | Two | Three | Four | Five | Six | Seven | Adults |  |  |
| *Didelphis* | 17/20 | 37/46 | 36/31 | 25/39 | 36/36 | 39/31 | 30/41 | 34/43 | 103/115 |  |  |
| *Monodelphis* (D) |  |  | 15/18 | 29/25 | 18/20 | 15/14 | 16/17 | 10/16 | 41/47 |  |  |
| *Sapajus* |  | 05/13 | 08/09 | 15/16 |  | 10/21 | 57/70 |  | 57/70 |  |  |
| Birth age class |  |  |  |  |  |  |  |  |  |  |  |
|  | 20 | 30 | 40 | 50 | 70 | 100 | 150 | 200 | 300 | 400 | Adults |
| *Monodelphis* (B) | 13/08 |  | 09/08 |  | 06/09 |  | 12/12 |  | 08/05 | 10/09 | 18/14 |
| *Calomys* | 14/07 | 14/11 |  | 25/27 |  | 35/25 |  | 29/18 | 16/14 | 12/19 | 92/76 |

**Figure A1:**
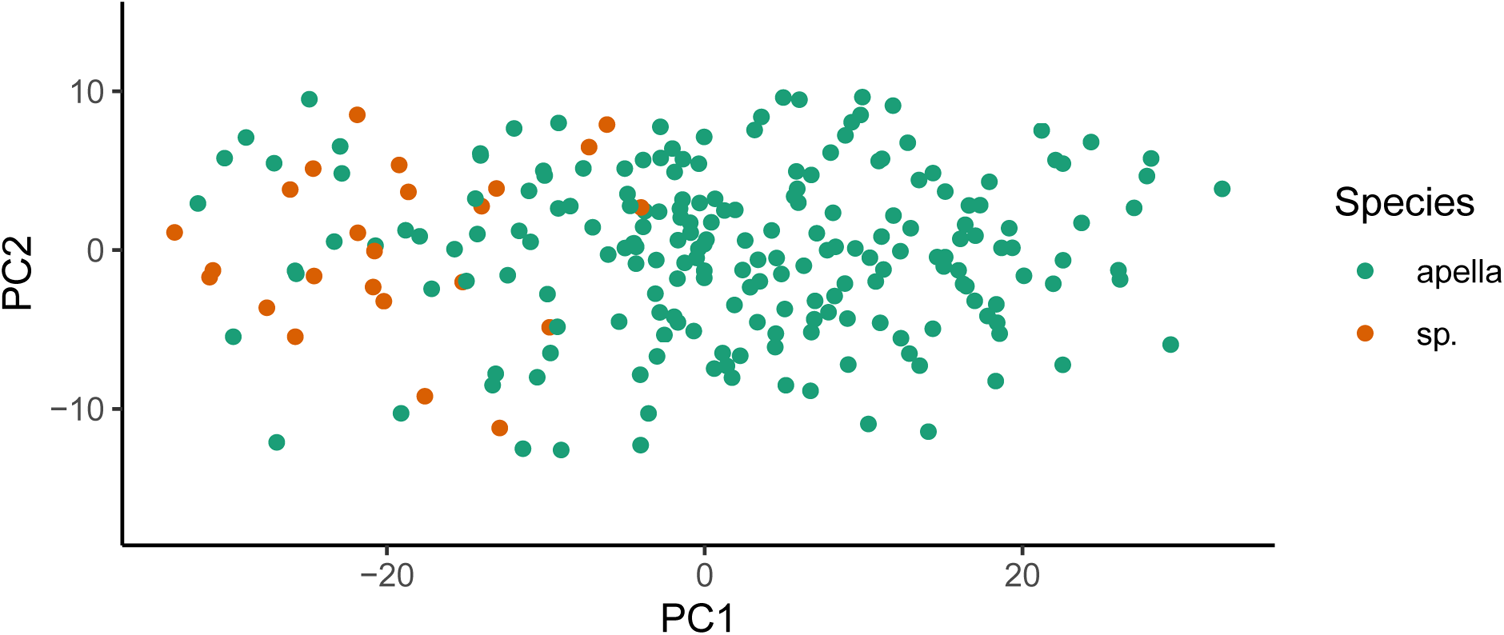
Principal component analysis for the *Sapajus* dataset, including specimens with unknown species.

**Figure A2:**
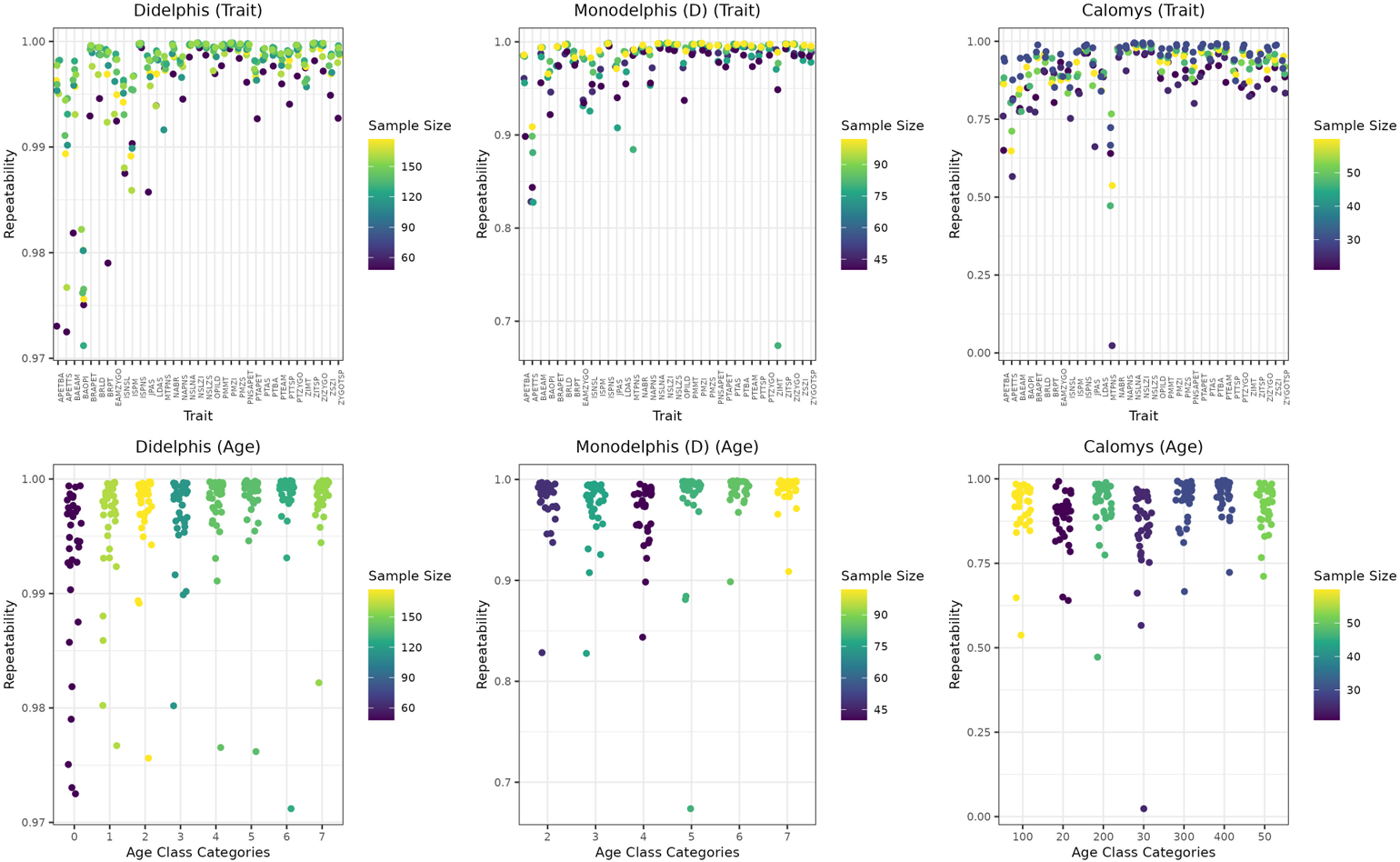
Repeatabilities for all measured traits for *Didelphis*, *Monodelphis* (D) and *Calomys* separated by trait, and age classes.

**Figure A3:**
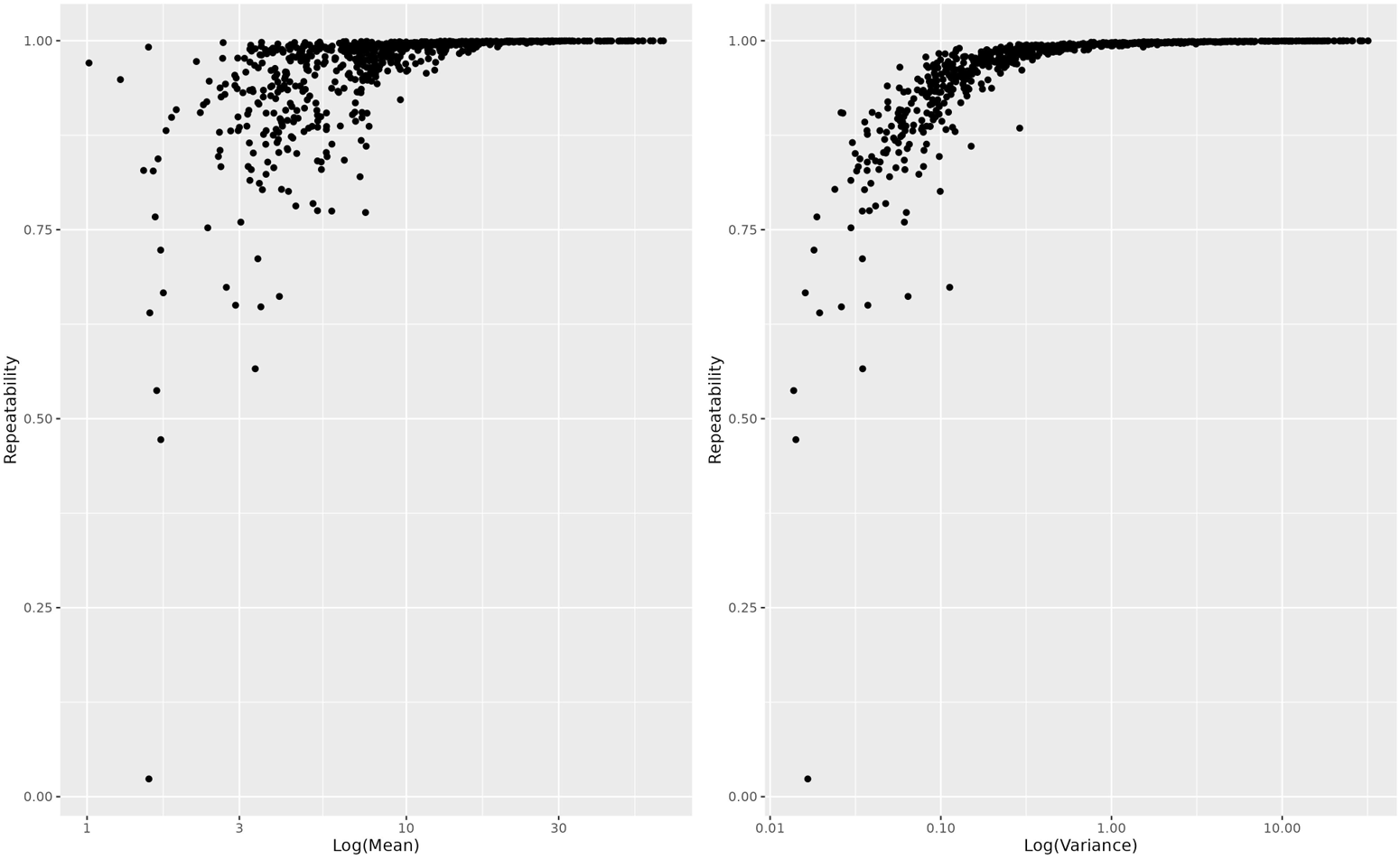
Scatterplot between repeatabilities for all traits, age classes, and ontogenetic series and respective traits’ mean (left) and variance (right).

**Figure A4:**
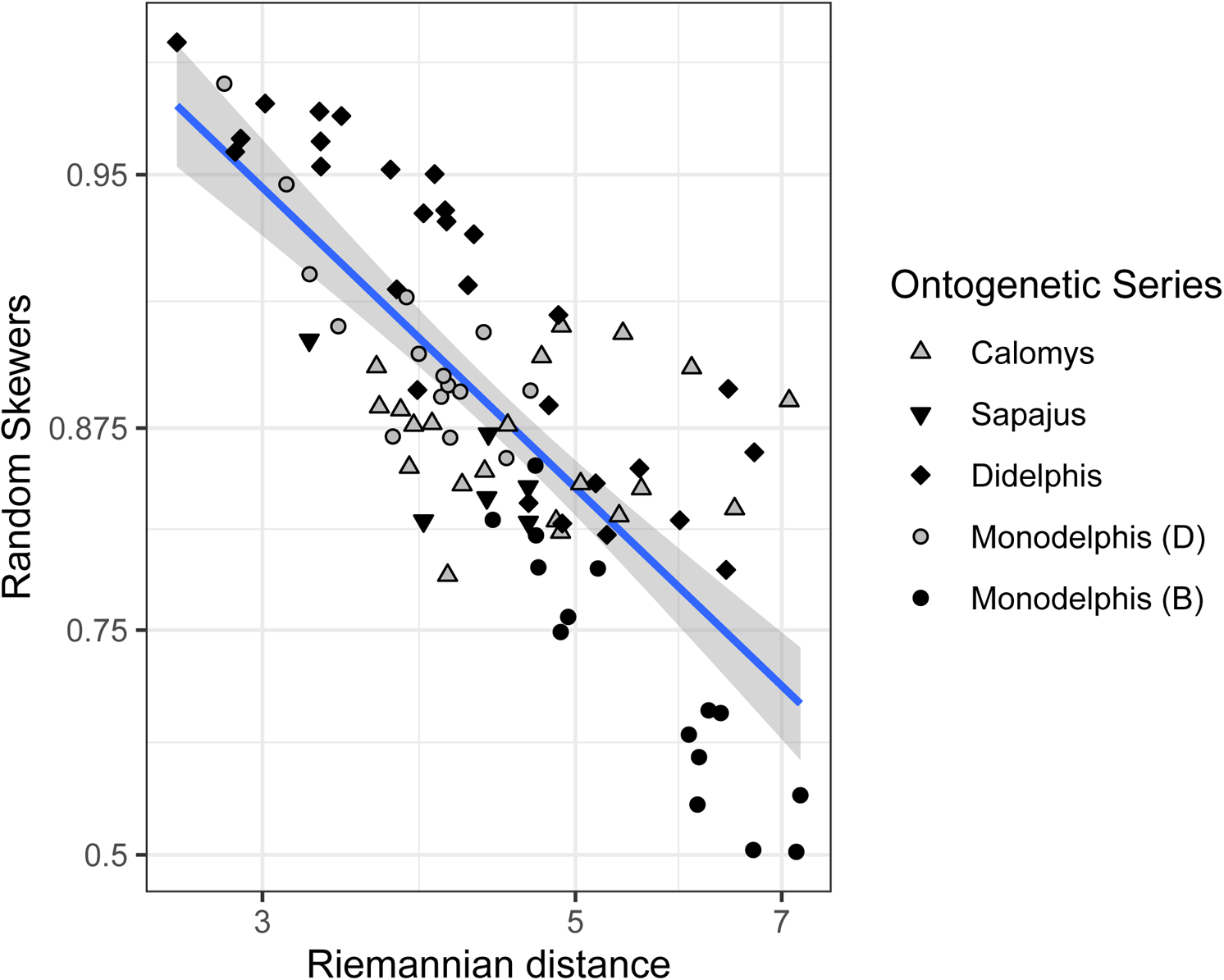
Relationship between Riemannian distance and uncorrected Random Skewers between age classes within the ontogenetic series.

**Figure A5:**
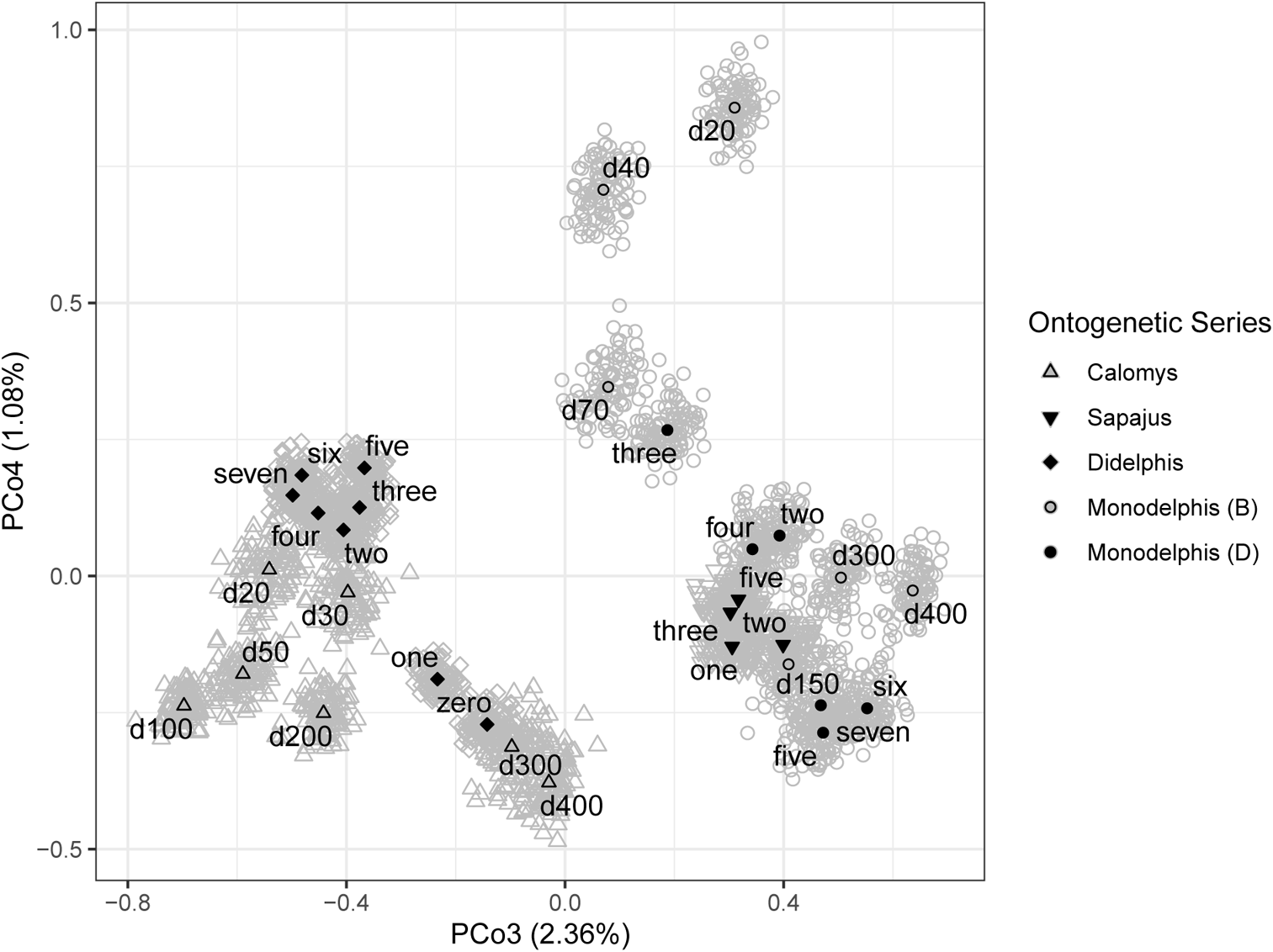
Distribution of age-specific **P**-matrices on the third and fourth principal coordinates based on the Riemannian distance. Black symbols represent the median of each posterior distribution of age-specific **P**-matrices within each ontogenetic series. Gray symbols represent 100 matrices from the posterior distribution of age-specific **P**-matrices within each ontogenetic series. The remaining PCoAs explained *<* 1% and are not figured.

**Figure A6:**
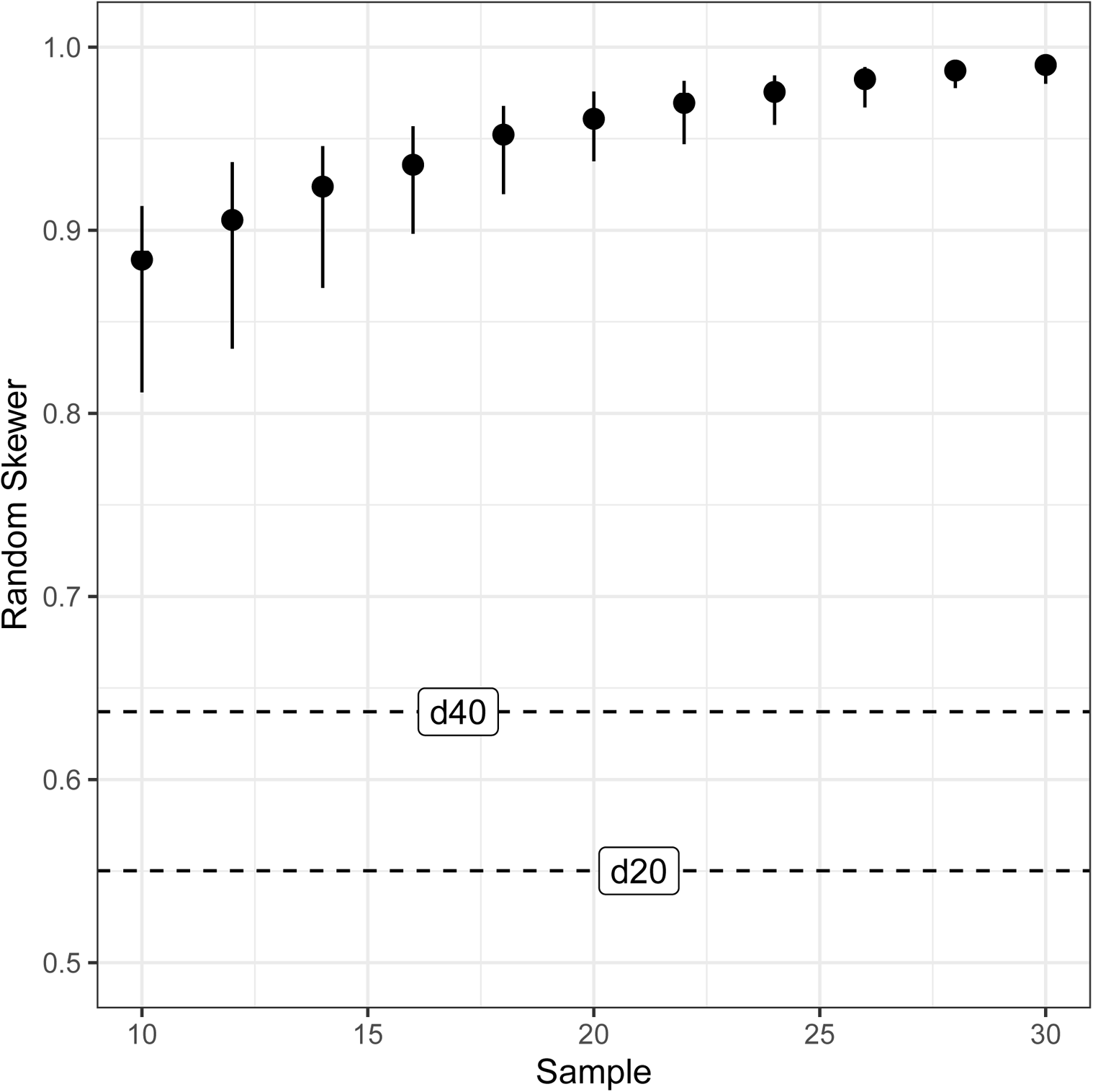
Rarefaction analysis of the *Monodelphis* (B) median of the posterior distribution for the adult **P**-matrix. Solid lines represent 95% confidence intervals for the Random Skewer statistics based on the resampling of the full *Monodelphis* (B) posterior median adult matrix using different sample sizes. Dashed lines represent the comparisons of the two posterior median younger age-classes (20 and 40) **P**-matrices against the posterior median adult **P**-matrix. If sample size was the main cause for differences in **P**-matrices patterns, we would expect that the Random Skewers value for **P**-matrices between birth age classes 20 or 40 and adult would fall within the confidence interval. Since this is not the case, at least part of the observed results are not sampling artifacts.

**Figure A7:**
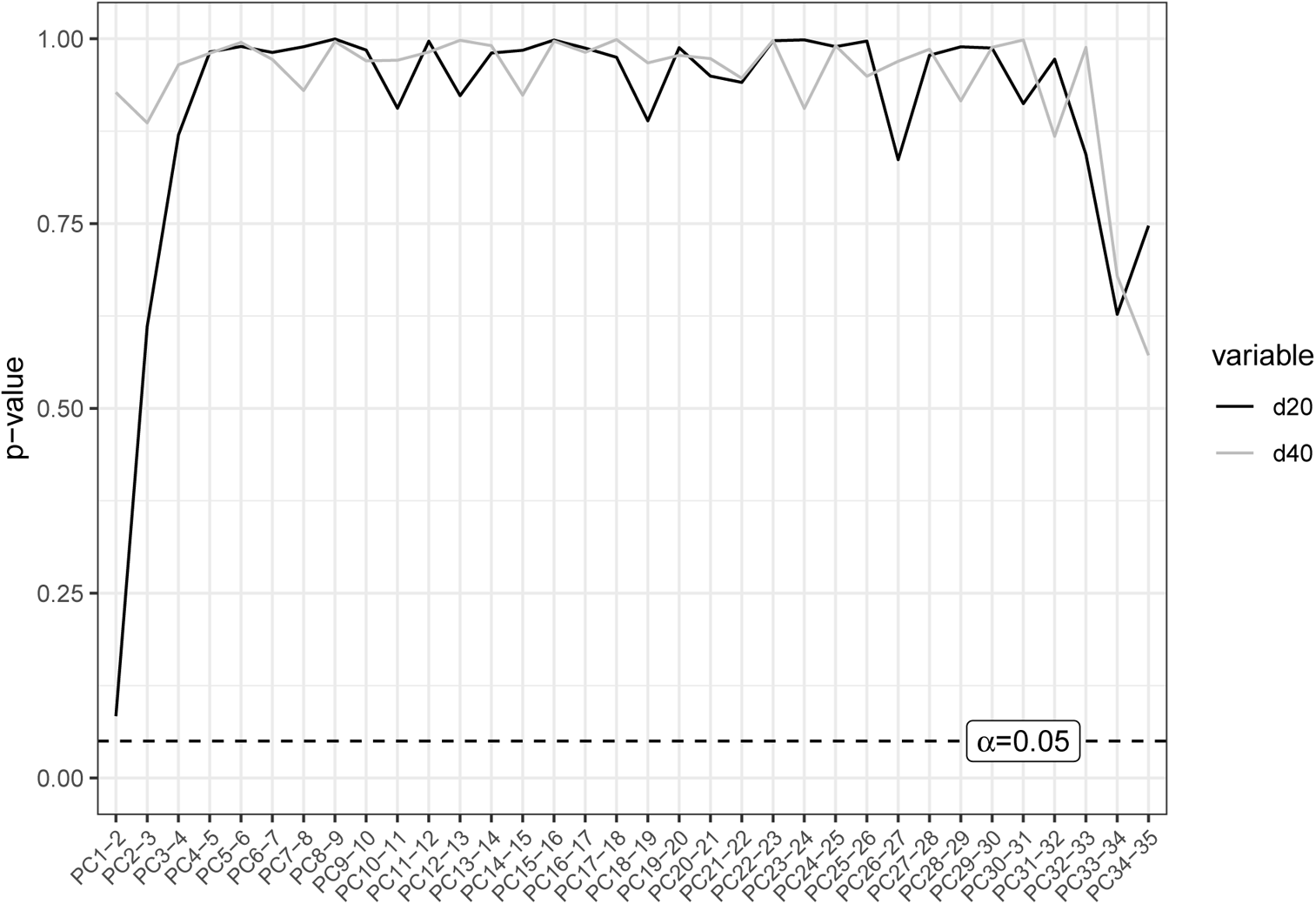
P-values for comparison of sequential eigenvectors in a relative eigenanalysis of the posterior median d20 and d40 **P**-matrices against the posterior median adult **P**-matrix for *Monodelphis* (B). The relative eigenanalysis is calculated as the eigendecomposition of the 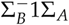, where Σ*_A_* is a covariance matrix of interest (the younger age-class matrices) and Σ*_B_* is a target matrix (the adult matrix in this case) (Bookstein and Mitteroecker, 2014). The eigenvectors of 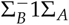 are the linear combination of traits that most differ between the two matrices. The leading principal componets (PCs) are the directions in which Σ*_A_* contains more variation than Σ*_B_*, and the last PCs are the ones in which Σ*_A_* contains less variation than Σ*_B_*. The middle PCs are thought to be the ones in which Σ*_A_* and Σ*_B_* are most similar. We employed a test to evaluate if sequential eigenvalues where significantly different from each other (Le Maître and Mitteroecker, 2019), and the p-values are displayed above. No PC was considered clearly divergent from the following one, hindering their interpretation.

**Figure A8:**
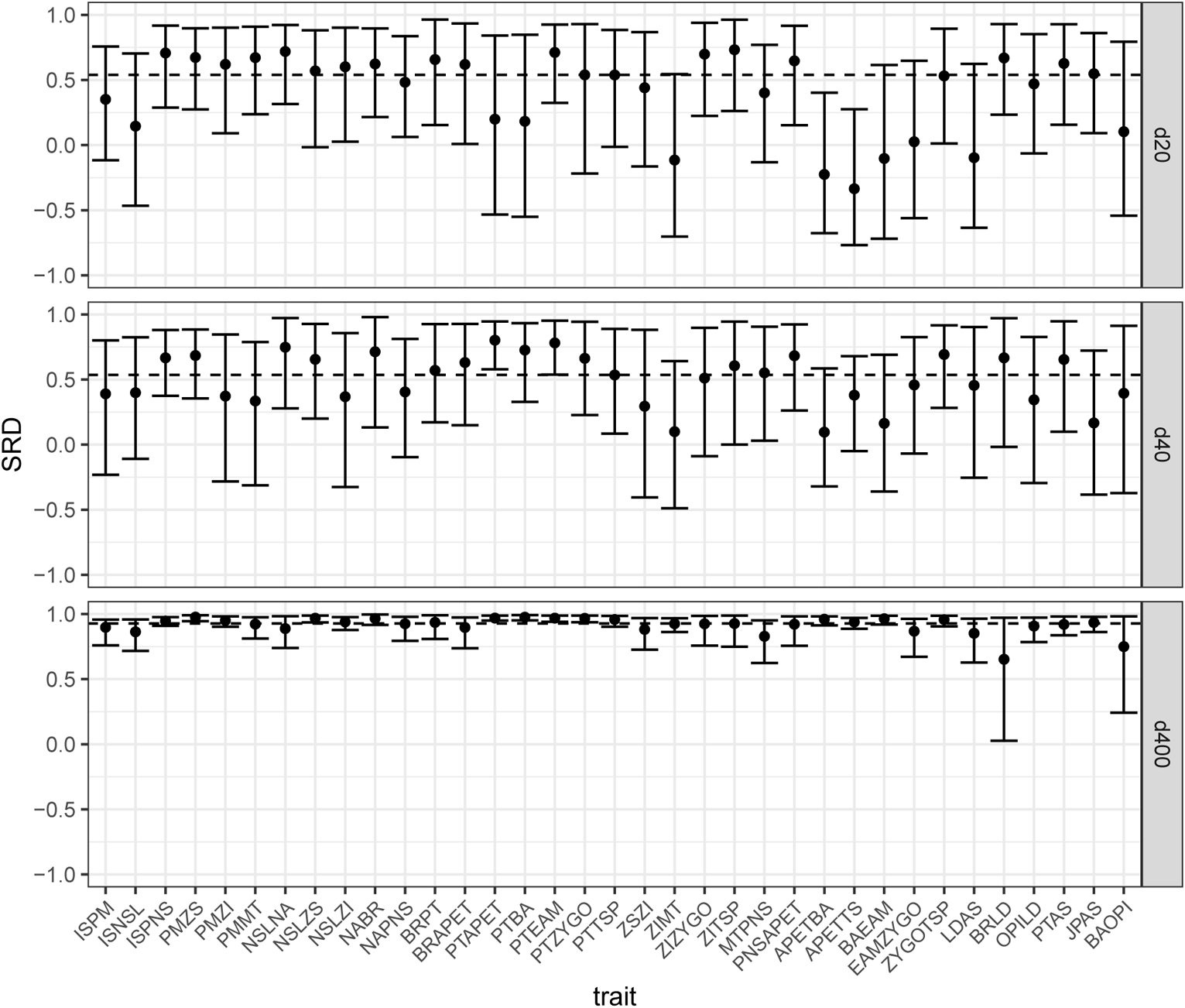
Selection response decomposition analysis (SRD) comparing posterior distribution median early (d20, d40) and late (d400) developmental stage **P**-matrices to the adult posterior median **P**-matrix for *Monodelphis* (B). SRD decomposes the Random Skewers equation into direct and indirect effects (Marroig et al., 2011). Traits with higher SRD values are responding similarly among matrices, while lower values mean that traits are responding differently. Confidence intervals are generated through 1000 random vectors, and the dashed line is the posterior median SRD value (a value similar to the uncorrected RS value). The SRD values for the d400 age class are given for comparison, showing the SRD profile for similar matrices. Both d20 and d40 differ from the adult matrix, but the dissimilarity do not seem to be restricted to a specific set of traits.

**Figure A9:**
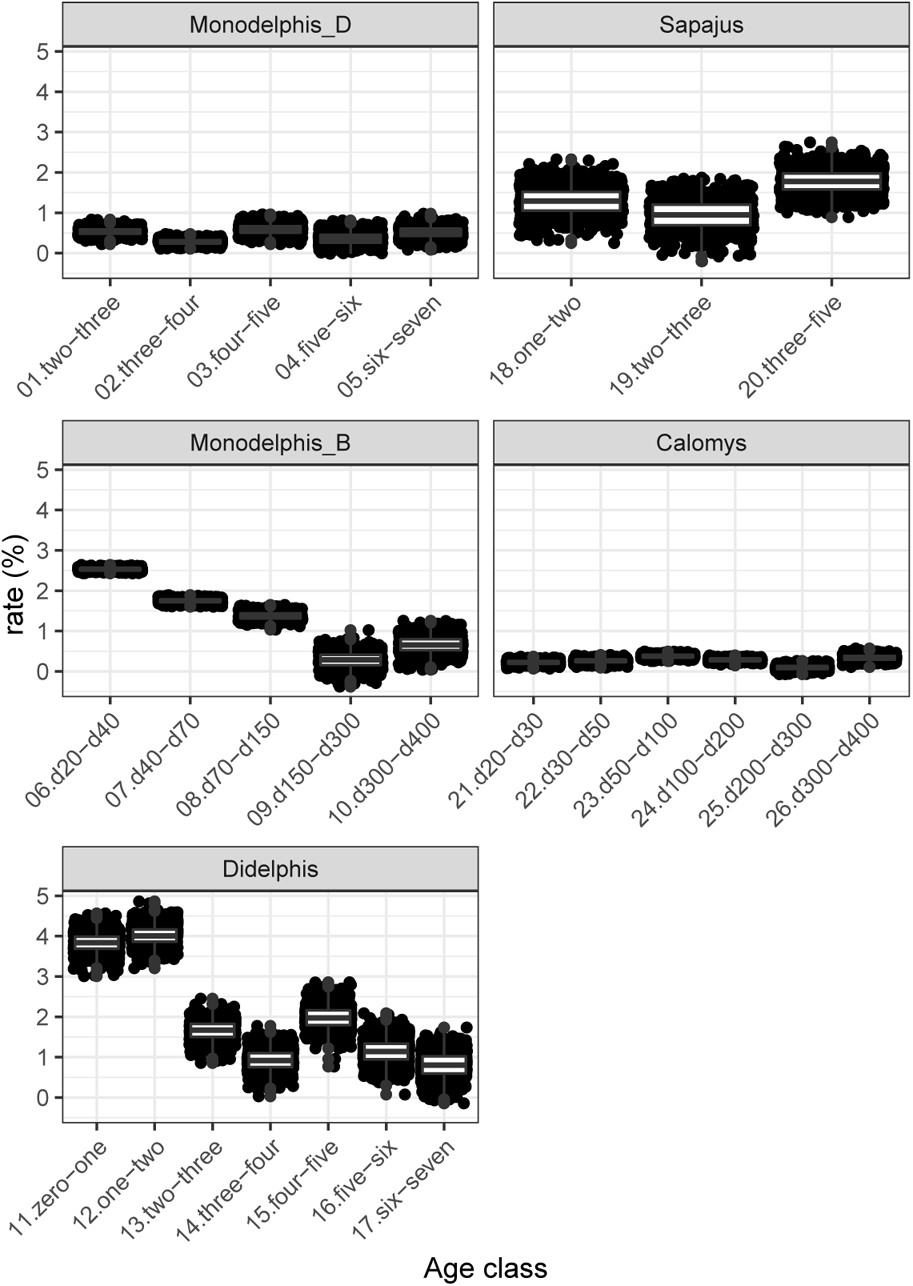
Growth rate for each ontogenetic series between contiguous age classes. Growth rate was calculated as the percentage increase in the average geometric mean of all cranial traits one age to the next. Distributions were generated by a Monte-Carlo resampling (1,000 times). Note that *Monodelphis* (B) and *Didelphis* younger age classes are growing faster than the other age classes. The fact that covariance patterns for *Monodelphis* (B) during lactation show a different pattern than older age classes while *Didelphis* do not, may be explained by differences in sampling specimens based on birth age classes and dental age classes, respectively. Dental age classes combine in a single age class specimens with different absolute ages (Richtsmeier et al., 1993; van Nievelt and Smith, 2005). This may inflate the total amount of morphological variance explained by size, particularly for dental age classes in which specimens are experiencing higher growth rates, since relatively small variation in absolute ages within the dental age class will result in relatively large variance in size. Since size is a major component of variance in Didelphimorphia skull traits, (Shirai and Marroig, 2010), the overestimation of size influence in younger dental age classes will lead to more similar covariance patterns between them and the older ones.

**Figure A10:**
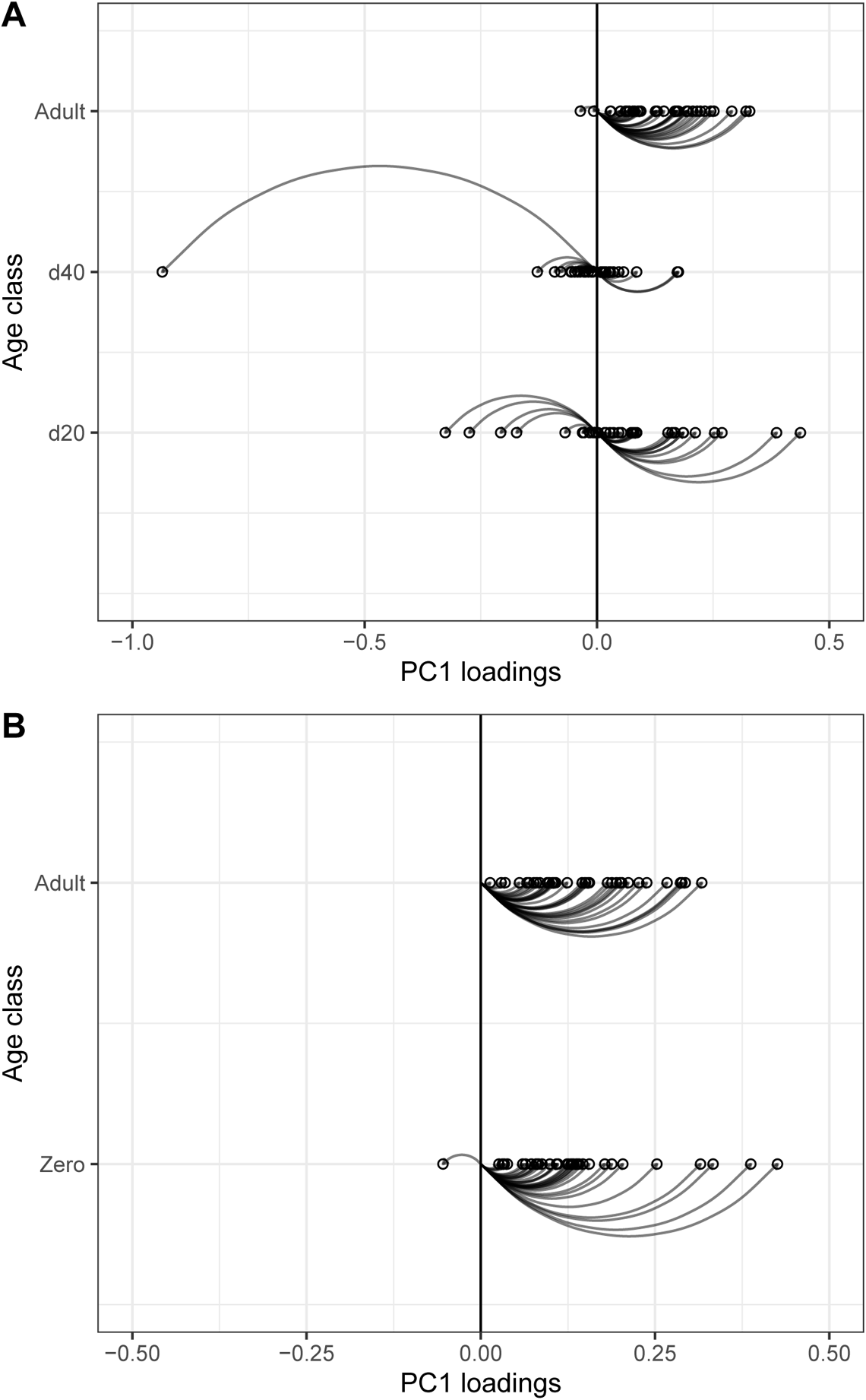
Loadings for the principal component 1 for both median of the posterior distribution for adults and the younger age-classes **P**-matrices for *Monodelphis* (B) (**A**) and *Didelphis* (**B**).

